# Orthogonal Drug Pooling Enhances Phenotype-based Discovery of Ocular Anti-angiogenic Drugs in Zebrafish Larvae

**DOI:** 10.1101/559781

**Authors:** Nils Ohnesorge, Temitope Sasore, Daniel Hillary, Yolanda Alvarez, Michelle Carey, Breandán N. Kennedy

**Author notes:** Correspondence: Prof. Breandán Kennedy.

## Abstract

Unbiased screening of large randomized chemical libraries *in vivo* is a powerful tool to find new drugs and targets. However, forward chemical screens in zebrafish can be time consuming and usually >99% of test compounds have no significant effect on the desired phenotype. Here, we sought to find bioactive drugs more efficiently and to comply with the 3R principles of replacement, reduction and refinement of animals in research. We investigated if pooling of drugs to simultaneously test 8-10 compounds in zebrafish larvae can increase the screening efficiency of an established assay that identifies drugs inhibiting developmental angiogenesis in the eye. In a phenotype-based screen, we tested 1760 small molecule compounds from the ChemBridge DIVERSet™ chemical library for their ability to inhibit the formation of distinct primary hyaloid vessels in the eye. Applying orthogonal pooling of the chemical library, we treated zebrafish embryos from 3 to 5 days post fertilization with pools of 8 or 10 compounds at 10 μM each. This reduced the number of tests from 1760 to 396. In 63% of cases, treatment showed sub-threshold effects of <40% reduction of primary hyaloid vessels. From 18 pool hits, we identified 8 compounds that reduce hyaloid vessels in the larval zebrafish eye by at least 40%. Compound 4-[4-(1H-benzimidazol-2-yl)phenoxy]aniline ranked as the most promising candidate with reproducible and dose-dependent effects. To our knowledge, this is the first report of a self-deconvoluting matrix strategy applied to drug screening in zebrafish. We conclude that the orthogonal drug pooling strategy is a cost-effective, time-saving and unbiased approach to discover novel inhibitors of developmental angiogenesis in the eye. Ultimately, this approach may identify new drugs or targets to mitigate disease caused by pathological angiogenesis in the eye *e.g.* diabetic retinopathy or age-related macular degeneration wherein blood vessel growth and leaky vessels lead to vision impairment or clinical blindness.

## Introduction

A decline in the number of patented novel chemical entities (NCEs) highlights the need for alternative drug discovery approaches (Scannell et al., 2012). Target-based drug discovery or *reverse pharmacology* has dominated recent decades. However, the development of more efficient, higher-throughput target-based approaches did not stall this decline (Rai and Sherkow, 2016). A renaissance is occurring in the use of phenotype-based drug discovery or *forward pharmacology*, an alternative approach which identifies drugs that change the observable traits of cells or organisms (Swinney and Anthony, 2011). Although generally not as efficient as target-based drug discovery, this approach enables unbiased or target-agnostic screening, thereby identifying unanticipated drugs and targets that modulate physiological or pathological phenotypes (Wang et al., 2010; Rennekamp et al., 2016). Furthermore, when performed in whole organisms the pharmacodynamics, pharmacokinetics and toxicology of a drug is evaluated in a complex physiological system (MacRae and Peterson, 2015; Wiley et al., 2017).

Zebrafish are a cost-effective vertebrate model for phenotype-based drug screens (Kitambi et al., 2009; Williams and Hong, 2016). Their small size, high fecundity and large clutches of transparent embryos, coupled with rapid and external development consolidates zebrafish as a robust model organism for drug discovery (North et al., 2007; Rezzola et al., 2016). As vertebrates, zebrafish have significant similarity to humans including orthologs of 80% of the expressed genome, and ability to investigate features of human physiology and disease (*e.g.* haematopoiesis, tissue regeneration, cancer and blindness), therefore providing a robust translational model (Wang et al., 2010; Li et al., 2015). Phenotype-based readouts include assays of development, behavior, metabolism and angiogenesis’(Peterson et al., 2000; Baraban et al., 2013; Rennekamp and Peterson, 2015) often applying bespoke reporter or mutant lines. In screens related to angiogenesis, the Tg(*fli1a*:GFP) or Tg(*flk1*:GFP) transgenic lines expressing GFP in all vascular endothelial cells allow facile visualization of vasculature development in specific organs (Lawson and Weinstein, 2002; Alvarez et al., 2007; Hartsock et al., 2014).

Aberrant ocular angiogenesis in diabetic retinopathy (DR) or age-related macular degeneration (AMD) leading to blindness is a major socio-economic problem (Bourne et al., 2013; Cunha-Vaz, 2014). Leaky vessels in the eye cause vitreous hemorrhage, retinal detachment and macular edema, leading eventually to complete loss of vision. The most successful therapies for these vasculopathies are antibodies targeting vascular endothelial growth factor (VEGF) administered directly into the eye (Querques et al., 2015). However, the intraocular injections are expensive and require repeat administration every 4-8 weeks. In addition, the injections are uncomfortable, increase risk of eye infection and ∼50% of patients become non-responsive, probably due to VEGF-independent resistance (Kwong and Mohamed, 2014). Thus, small molecule organic chemicals exerting anti-angiogenic properties via alternative, additive or synergistic pathways offer potential to be developed as novel stand-alone or combinatorial drugs.

To date, the majority of phenotype-based drug screens in zebrafish test chemical libraries of 1000-5000 compounds (Rennekamp and Peterson, 2015). Automated or robotic technology for zebrafish sorting and drug treatment facilitates higher-throughput, which then becomes rate-limited by the time for analysis (Burns et al., 2005; Vogt et al., 2009; Wheeler and Brandli, 2009; Graf et al., 2011; Breitwieser et al., 2018). As a complementary approach, we applied a “drug pooling” method to enable faster identification of the most promising compounds. In contrast to the “one compound, one well” approach of previous screens, in “drug pooling” combinations of several compounds are tested first in a primary screen and potential hits are confirmed in secondary screens. The rationale behind this approach is that in randomized chemical libraries, only a small fraction (0.6-1.7%) are bioactive compounds and most substances can be quickly identified as inactive via negative results of a tested pool (Kainkaryam and Woolf, 2009; Clifton et al., 2010; Peal et al., 2011; Reynolds et al., 2016; Saydmohammed et al., 2018). By, effectively lowering sample numbers when investigating large chemical libraries this accelerates scientific findings and conducts drug screening in accordance with the 3R principles.

## Material and Methods

### Zebrafish husbandry

The zebrafish transgenic line Tg(*fli1a*:eGFP)^y1^ that expresses enhanced green fluorescent protein (GFP) under the control of an endothelial specific promoter was raised under standard conditions (28°C, pH 7.2-7.4, 14/10 hrs light/dark cycle) with pH and dissolved oxygen routinely monitored (Lawson and Weinstein, 2002; Goodwin et al., 2016). The adult fish were fed twice daily with live ZM Ltd artemia and Lillico zebrafish diet. Zebrafish eggs were collected in 10 cm petri dishes in Hanks embryo medium (Westerfield, 1995). Embryos were staged and normal development was confirmed under an Olympus stereo microscope (SZX10). All experiments were approved by UCD ethics committee and conducted according to EU directive 2010/63/EU on the protection of animals used for scientific purposes.

### Compounds tested

The ChemBridge DIVERSet™ library contains compounds with drug like properties, designed with MW ≤500, clogP ≤5, tPSA ≤100, rotatable bonds ≤8, hydrogen bond acceptors ≤10 and hydrogen bond donors ≤5 and an absence of non-drug like chemical groups. 1760 random small organic molecules of the ChemBridge DIVERSet™ library dissolved in 100% DMSO at a stock concentration of 10 mM were screened as attenuators of zebrafish ocular vasculature development.

### Drug pooling

The DIVERSet™ library compounds were provided as 80 compounds per 96 well plate. To increase screening throughput, 10 compounds of a row or 8 compounds of a column were orthogonally pooled and tested in final concentration of 10 μM per compound in 1% DMSO. The 80 compounds of each plate were thus tested in 18 pools with every compound represented in two different pools (***Fig. 1B***).

**Figure 1:**
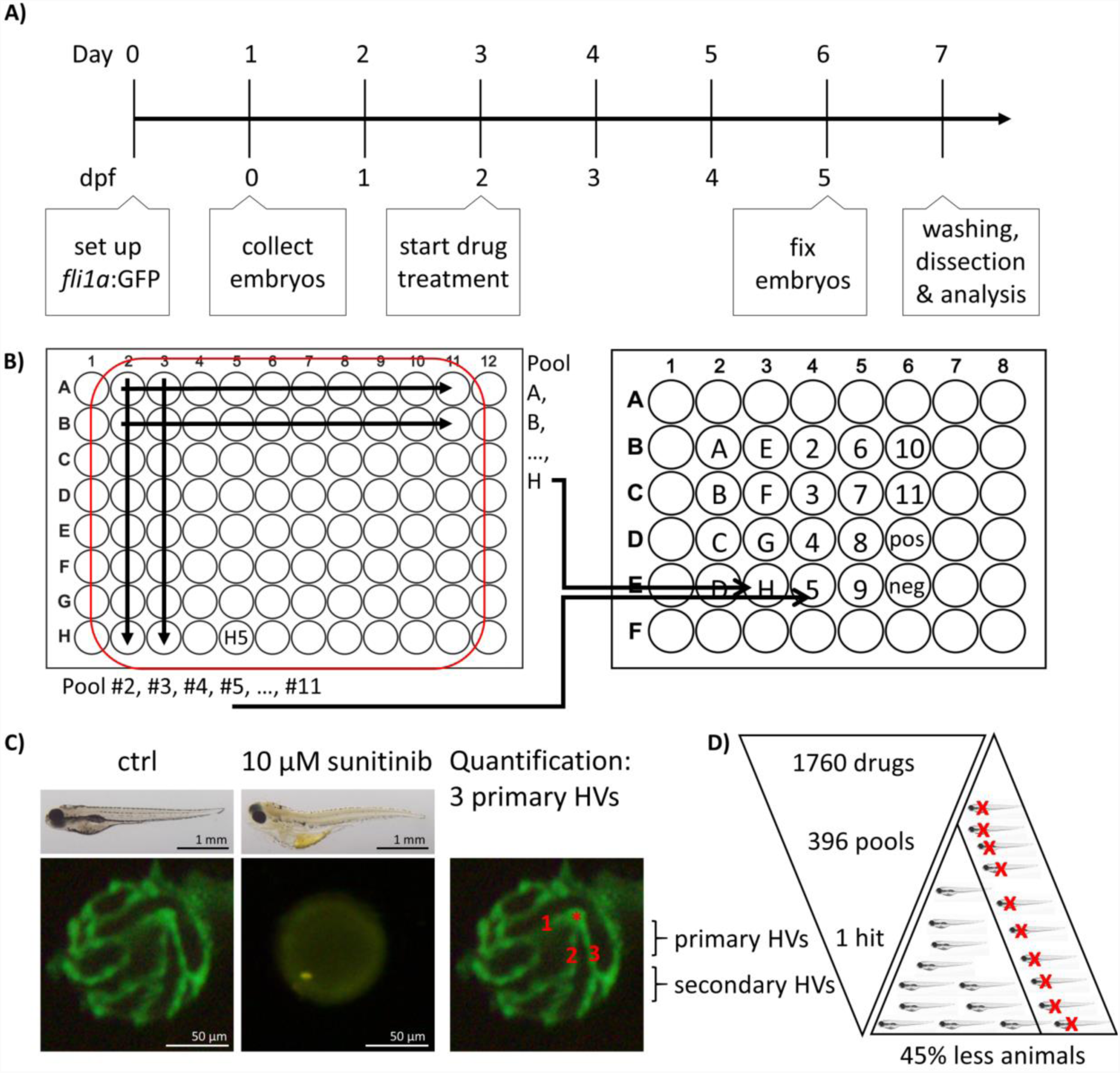
Method overview for orthogonal drug pooling. (A) Overview of drug treatment protocol. Tg(*fli1a*:eGFP) positive embryos were treated from 2-5 dpf and screened for intraocular vascular defects to assess the anti-angiogenic potential of test chemicals. Sunitinib was used as positive anti-angiogenic control and 1% DMSO as negative vehicle control. (B) In each 96 well plate, the 8 compounds of 10 columns and the 10 compounds of 8 rows were assembled in pools (left panel). This orthogonal pooling protocol reduced 80 individual compounds to 18 test pools. Every compound is represented in two pools (right panel). (C) The primary hyaloid vessel assay readout assesses the lenses dissected (lower left and middle box) from fixed larvae and quantification of the number of primary hyaloid vessels emerging from the optic disk (asterisk) on the back of the lens counted manually under a stereomicroscope (lower right box). (D) In total, 1760 compounds were analyzed, combined in 396 pools, resulting in one confirmed hit using this method. This assay replaces animal use with immature larval forms and the orthogonal pooling reduces the number of immature larvae needed by 45%.

### Drug treatments & Quantification

To determine the effect of test drugs on hyaloid vessel (HV) formation, 5 embryos were treated at 2 days post fertilization (dpf) with single or pooled compounds at a starting concentration of 10 μM in a volume of 400 μL embryo medium (***Fig. 1A***)(Alvarez et al., 2007; Hartsock et al., 2014). At 5 dpf, embryos were fixed with 4% PFA and lenses dissected from the eye to count connected primary hyaloid vessels (***Fig. 1C***) (Alvarez et al., 2007). To assess inhibition of angiogenesis in the intersegmental vessel (ISV), embryos were treated from 6 to 72 hours post fertilization (hpf). At 1.5 dpf, the ISVs that sprout from the dorsal aorta and elongated dorsally have reached the most dorsal region of the trunk and formed a T shape. Once these vessels connect they form a pair of dorsal longitudinal anastomotic vessels (DLAV) (Isogai et al., 2001). For quantification, all ISVs that reached the most dorsal position and were connected to the DLAV were manually counted as 1. Absent vessels or sprouts without connection to the DLAV were counted as 0. Dead larvae were not considered for quantification. In wells with 3 or more dead larvae, the combination of drugs was considered “toxic”. Unless all compounds of a toxic combination were represented in other non-toxic pools, they were tested again individually. 1% DMSO was used for negative vehicle control in pools and 0.1% DMSO as vehicle control for individual treatments. 10 μM sunitinib (Sigma Aldrich #PZ0012) was used as positive control (Faivre et al., 2007; Hao and Sadek, 2016). The number of blood vessels was determined using an Olympus stereo microscope (SZX10) and representative pictures taken using Olympus DP71 camera and CellSens Standard software (***Fig. 1C***).

### Statistical Analysis

We wish to assess whether the means of two groups are statistically different from each other. An independent sample t-test will evaluate if the data supports this claim but requires that the population of the sample is approximately normally distributed within each group and the population variances of the two groups are equal. We use a Shapiro-Wilk test to assess whether the sample is drawn from a population with a normal distribution and a Fligner-Killeen test to determine if the variances of the two populations are equal. If the Shapiro-Wilk test and the Fligner-Killeen test hypotheses are not rejected, then an independent sample t-test is appropriate. If only the hypothesis of the Fligner-Killeen test is rejected, a t-test adjusted for unequal variance (Welch’s t-test) is appropriate. If the hypothesis of the Shapiro-Wilk test is rejected, then a Mann-Whitney U test is suitable. For each test if the p-value is less than 0.05 we reject the null hypothesis and conclude there is evidence of a significant difference between the groups (* p-value <0.05, ** p-value <0.01 and *** p-value <0.001).

## Results

### Orthogonal Drug Pooling

We decided on an orthogonal pooling strategy to evaluate both the feasibility and benefits of drug pooling for identification of library chemicals which attenuate hyaloid vessel development (***Fig. 1***). Our pooling strategy combined 10 drugs of each plate row (*e.g.* pool H) and 8 drugs from each plate column (*e.g.* pool 5) into single pools (***Fig. 1B***) with a final concentration of 1% DMSO. This ensured that every compound was present in at least two pools (*e.g.* H5 in pool H and pool 5) making it easier to distinguish true positives. As a phenotype-based readout for anti-angiogenic activity, we counted the number of primary hyaloid vessels in lenses dissected from 5 dpf eyes of Tg(*fli1a*:eGFP) embryos, following 3 days of drug treatment (***Fig. 1C***). Sunitinib, a potent multi-tyrosine kinase inhibitor with known anti-angiogenic properties at 10 μM was used as positive control. This method allows testing of 80 compounds from one 96-well plate in 18 pools or the library of 1760 compounds (22 plates) to be tested in 396 pools (***Fig. 1D***).

### Proof of Principle

We first determined if the final pool concentration of 1% DMSO may non-specifically generate an anti-angiogenic phenotype. In a dose-response analysis (ranging from 0.1 – 3%) DMSO, concentrations up to 2% DMSO had no effect on primary HV (PHV) formation (***Fig. 2A***). To implement and validate the adopted method, we first pooled 10 compounds of row D and 8 compounds of column 4 from randomly chosen Diverset® plate 30331 (***Fig. 2B***). Pool 4 (3.0 +/− 1.0 PHV) was indistinguishable from the vehicle controls (3.5 +/− 0.6 PHV) (***Fig. 2B***). Pool D induced a slight (2.6 +/− 0.5 PHV) but significant (p<0.05) reduction by 25% in hyaloid vessel number (***Fig. 2B***). None of the ten constituent compounds (D2-D11) from pool D exerted a significant anti-angiogenic effect on their own, however, D3 by itself reduced hyaloid vessel (2.8+/− 0.5 PHV OR 21% reduction) similar to pool D (***Fig. 2B***). In contrast, when compound D4 was replaced with 10 μM sunitinib in pool D or pool 4, robust antiangiogenic activity (0.2 +/− 0.4 PHV, 94% reduction) was observed, equivalent to treating with 10 μM sunitinib alone (0.3 +/− 0.5 PHV, 91% reduction). We conclude that active anti-angiogenic compounds can be detected in our drug pooling screen.

**Figure 2:**
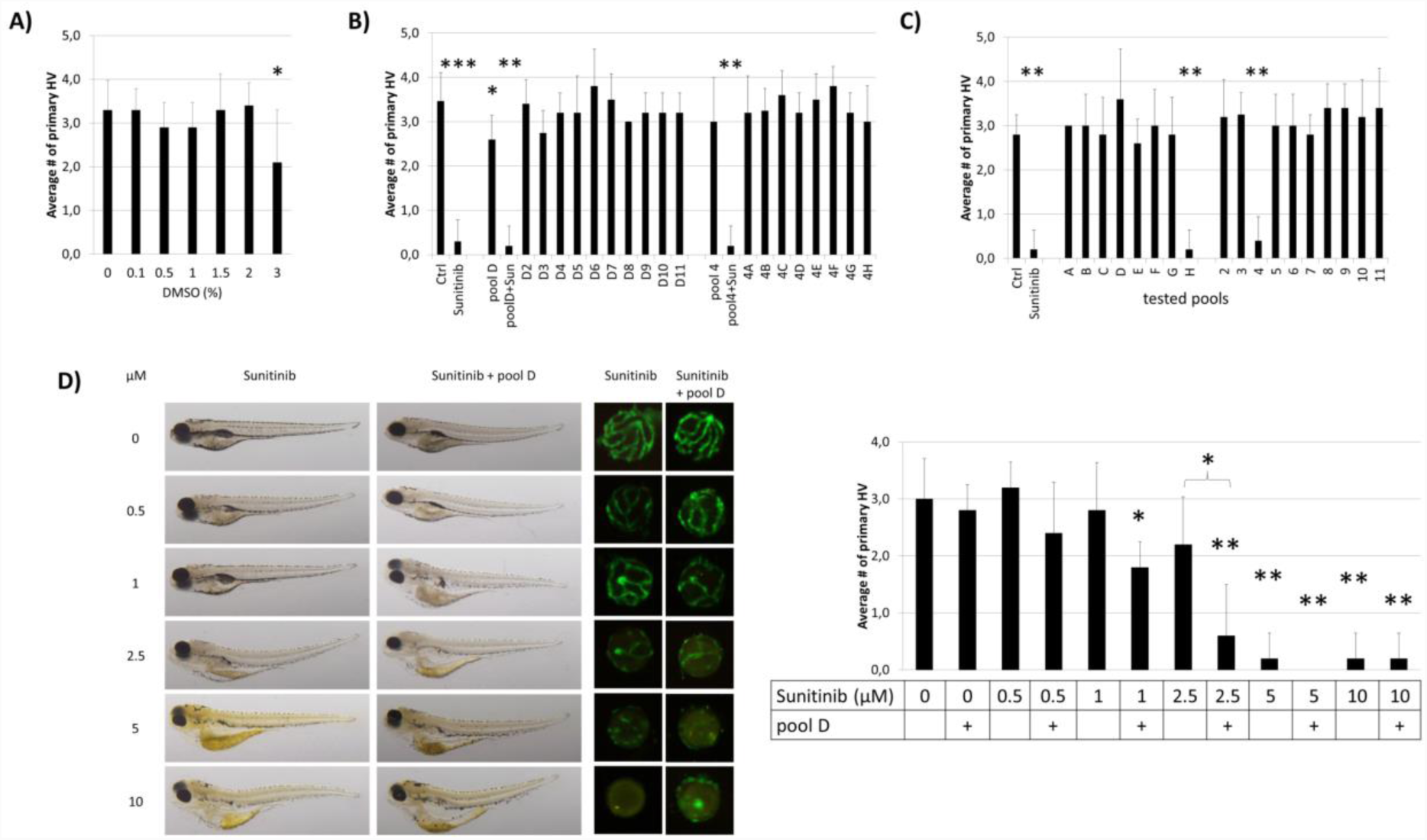
Proof-of-Principle. (A) Effect of DMSO concentration on HV assay. 1% DMSO vehicle has no effect on HV numbers. (B) The compounds of row D (D2-D11) and column 4 (4A-4H) in plate 30331 were tested individually and as pools. When 10 μM sunitinib was added to the pool “D” or “4” (pool D+Sun or pool4+Sun), its anti-angiogenic effects were as pronounced as if tested on its own (Sunitinib). (C) When compounds of plate 30251 were tested in pools, well H4 which was replaced with 10 μM sunitinib, was easily detected as anti-angiogenic in both pools (“H” & “4”) containing H4. (D) A dose-response analysis with different concentrations of sunitinib alone or added to pool D in plate 30331 proves that weaker anti-angiogenic effects can be reliably detected in pools. Shown are representative bright-field images of larvae, corresponding fluorescent images of dissected lenses and the quantification of primary hyaloid vessel numbers (right). Data shown are means +/− SD. Statistical significance was calculated by t-test and Mann Whitney test (* p < 0.05, ** p < 0.01, *** p < 0.001).

To scale-up the *proof-of-concept* screen, we tested all 80 compounds of randomly chosen Diverset® plate 30251 in 18 pools (***Fig. 2C***). Compound H4 was replaced with 10 μM sunitinib as a positive control. Reassuringly, pools H and 4, which contain 10 μM sunitinib, were true positives resulting in 0.2 +/− 0.4 and 0.4 +/− 0.5 primary hyaloid vessels reflecting a 93% and 87% reduction respectively. There was a clear efficacy difference between these pools and the non-spiked, inactive pools (***Fig. 2C***).

To address the concern that pooling compounds might mask active compounds, we performed a dose-response curve for sunitinib to determine if the concentration at which sunitinib no longer produced significant anti-angiogenic activity was different when administered alone compared to in D pools from Diverset® plate 30331 (***Fig. 2D***). Larvae treated with pool D alone were indistinguishable (2.8 +/− 0.4 PHV, p>0.6) from 1% DMSO vehicle controls. In general, the activity of sunitinib was equivalent in pools or alone, producing robust, significant anti-angiogenic activity at 5 and 10 μM concentrations (***Fig. 2D***). At 2.5 μM, sunitinib showed a significant reduction (p<0.05) in hyaloid vessel number in the pool and not with sunitinib alone (p>0.1), and the fold reduction was significantly different (0.6 +/− 0.9 and 2.2 +/− 0.8 PHV). This was also reflected in embryo morphology with oedema present when treated with 1 μM sunitinib in pool D but not with 1 μM sunitinib alone and more severe oedema forming at 2.5 μM sunitinib in pool D than with 2.5 μM sunitinib alone (***Fig. 2D***). In conclusion, our combinatorial approach was sufficient to identify anti-angiogenic compounds present in drug pools.

### Self-deconvoluting library screen

Theoretically every active compound should present in two pools (one row and one column). We additionally applied a cutoff of at least 40% reduction on primary hyaloid vessels number to be considered a hit. Smaller reductions were considered more likely to be false positives due to combinatorial effects. In the pooling screen, we tested 396 pools, representing 1760 compounds, for anti-angiogenic efficacy in the hyaloid vessel assay. Analysis of Diverset® plate #30328 depicts a representative screen result (***Fig. 3A***). In that plate, less than three larvae survived treatment in four pools (#30328 A, H, 4, 7). When compounds A4, A7, H4 and H7 were re-tested individually, none showed efficacy or toxicity (***Fig. 3B***). Notably, three of 18 pools (#30328 D, G & 9) showed a significant reduction in HV vessel numbers (***Fig. 3A***). Two pools exceeded our threshold of 40% PHV reduction (#30328G 75% reduction, p<0.05 and #30328-9 88% reduction, p<0.01) (***Fig. 3A***). To determine if a single compound present in pools G and 9 was responsible for the reduction in PHV, the 17 constituent compounds were tested individually. Compound G9 (chemical name: 4-(2,4-dichlorophenoxy)-N,N-diethyl-1-butanamine) initially reproduced significant anti-angiogenic activity but upon repeat testing in dose-response assays its activity-toxicity profile was highly variable and was eliminated from further follow up (***Fig. 3C-E***).

In contrast to plate #30328, treatment of larvae with pooled compounds of plate #30325 resulted in larval death of 50% and severe developmental problems in all remaining wells. Consequently, all 80 compounds of that plate were tested individually (***Fig. 3F***). Two compounds produced significant anti-angiogenic activity (7A, 7D with PHV 2.3+/− 0.5, 2.4+/− 0.5 and p-values 0.01 and 0.015, respectively), 3 resulted in a significant increase in primary hyaloid vessels (4G, 5H, 8B with PHV 3.8+/− 0.4, 3.8+/− 0.4, 4.2+/− 0.8 and all with p-values of 0.01) and 2 were toxic (10G, 11B). None of the significant changes was greater than our selected threshold of 40% and therefore not considered a hit.

**Figure 3:**
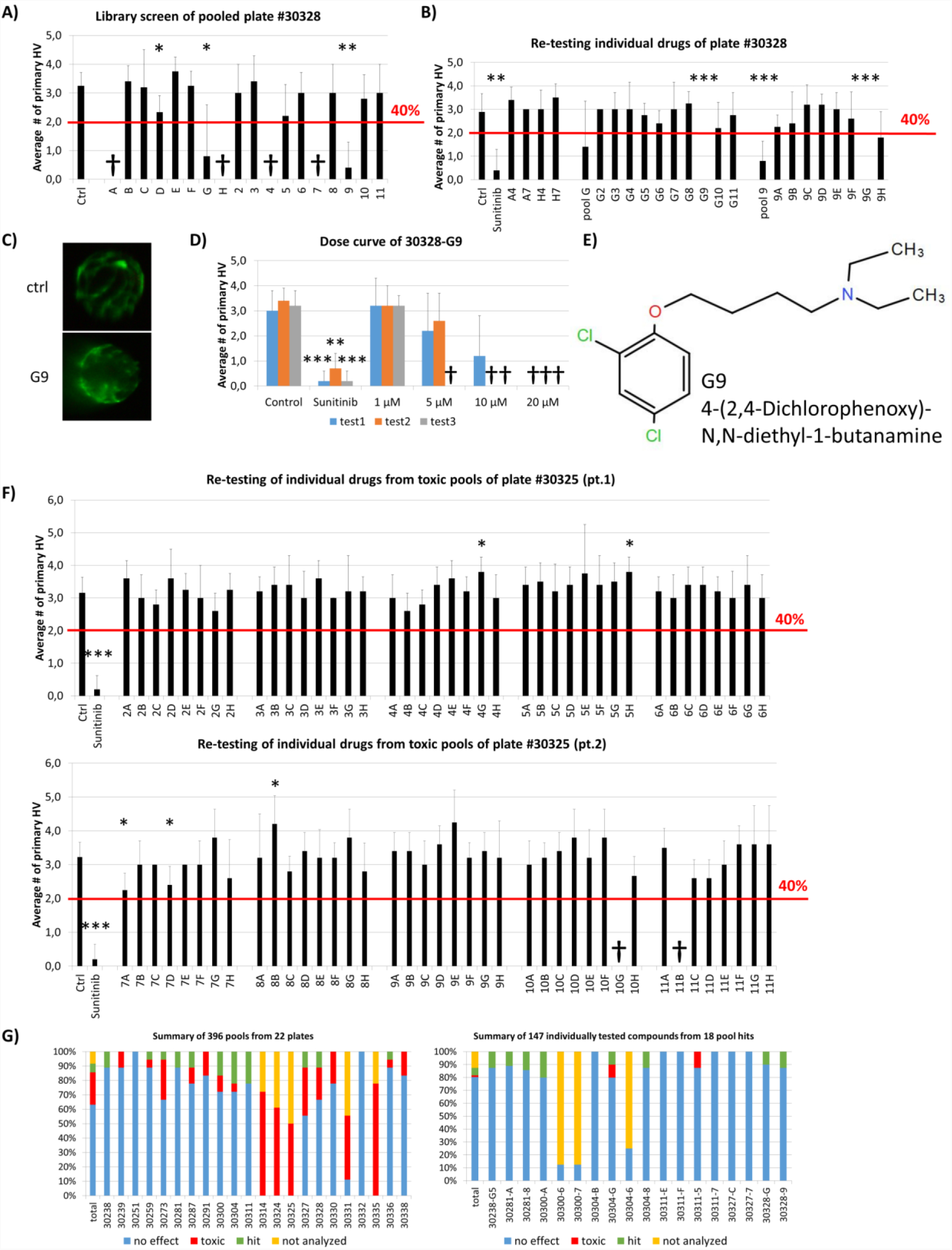
self-deconvoluting library screen. (A) The screen of plate #30328 is shown as a representative example where 80 drugs (A2-H11) were analyzed in 18 pools (A-H, 2-11). 4 pools were lethal to the embryos (A, H, 4, 7). Three pools showed a significant reduction in blood vessel numbers, but only two (G, 9) exceeded the threshold of 40% reduction in HV numbers. (B) Individually re-tested compounds from toxic pools or of pool hits from plate #30328 identified G9 as promising candidate for further testing with >40% PHV reduction. (C) Representative images from DMSO control or 10 μM G9 treated zebrafish lenses showing the GFP-positive hyaloid vessels. (D) Three independent dose curves (test1-3) of G9 drug treatments showed strong toxicity at concentrations of >5 μM and lack of reproducible anti-angiogenic activity. (E) The chemical structure of and chemical name (4-(2,4-Dichlorophenoxy)-N,N-diethyl-1-butanamine) of G9. (F) Individual re-testing of all 80 compounds from plate #30325 that were toxic in pools. No single compound reduced HV >40% and 2 were lethal at 10 μM (10G, 11B). (G) Summary of library screen. Pooled drug screens were conducted for 22 plates containing 1760 different compounds. In 63% of cases, combinations of 8 or 10 pooled drugs did not reduce HV numbers >40%. 22% of pools caused significant developmental defects or were lethal. Of the 147 compounds identified in 18 pool hits, 80% had no significant effect when tested individually, 2 were toxic and 8 were considered hits that significantly reduce hyaloid vessel number. Data in (A), (B), (D), (F) are means +/− SD. Statistical significance calculated by t-test and Mann Whitney test (* p < 0.05, ** p < 0.01, *** p < 0.001).

In total, of the 396 pools screened, 63% didn’t surmount the selected threshold of at least 40% PHV reduction compared to the control (***Fig. 3G***) and were dismissed. More problematic was the 89 or 22% toxic pools, wherein survival rates were under 50% (***Fig. 3G***). Notably, these predominantly originate from 5 plates with 55 toxic pools, and an additional 12 plates with 34 toxic pools (***Fig. 3G***). To test all of the constituent compounds would require testing 427 individual drugs. In practice, we tested 134 individual drugs from toxic pools and 7% of them showed toxicity and none exerted significant anti-angiogenic activity. In total, 24 pools reduced primary HV by greater than 40% (***Fig. 3G***). 6 of these had no corresponding hit in an orthogonal pool and were dismissed as false positives. The remaining 18 pools comprised of 147 compounds that were selected for individual testing (***Fig. 3B***, ***3G right panel***). In second round screens, 8 drug hits were identified but 7 of them did not exert a dose-dependent anti-angiogenic activity in tertiary screens.

Significantly, compound 30238-G5 showed anti-angiogenic activity when re-tested individually in secondary HV assays and also demonstrated a dose-dependent response at concentrations between 1 and 20 μM in tertiary screens (***Fig. 4A-C***). At 10 and 20 μM, G5 caused primary HV numbers to reduce by 25 or 50%, respectively, with no observed adverse effect on survival or overall morphology (***Fig. 4C left panel***). When tested in the alternative inter-segmental vessel (ISV) assay, G5 was inactive indicating a specific anti-angiogenic effect in the eye (***Fig. 4D***).

**Figure 4:**
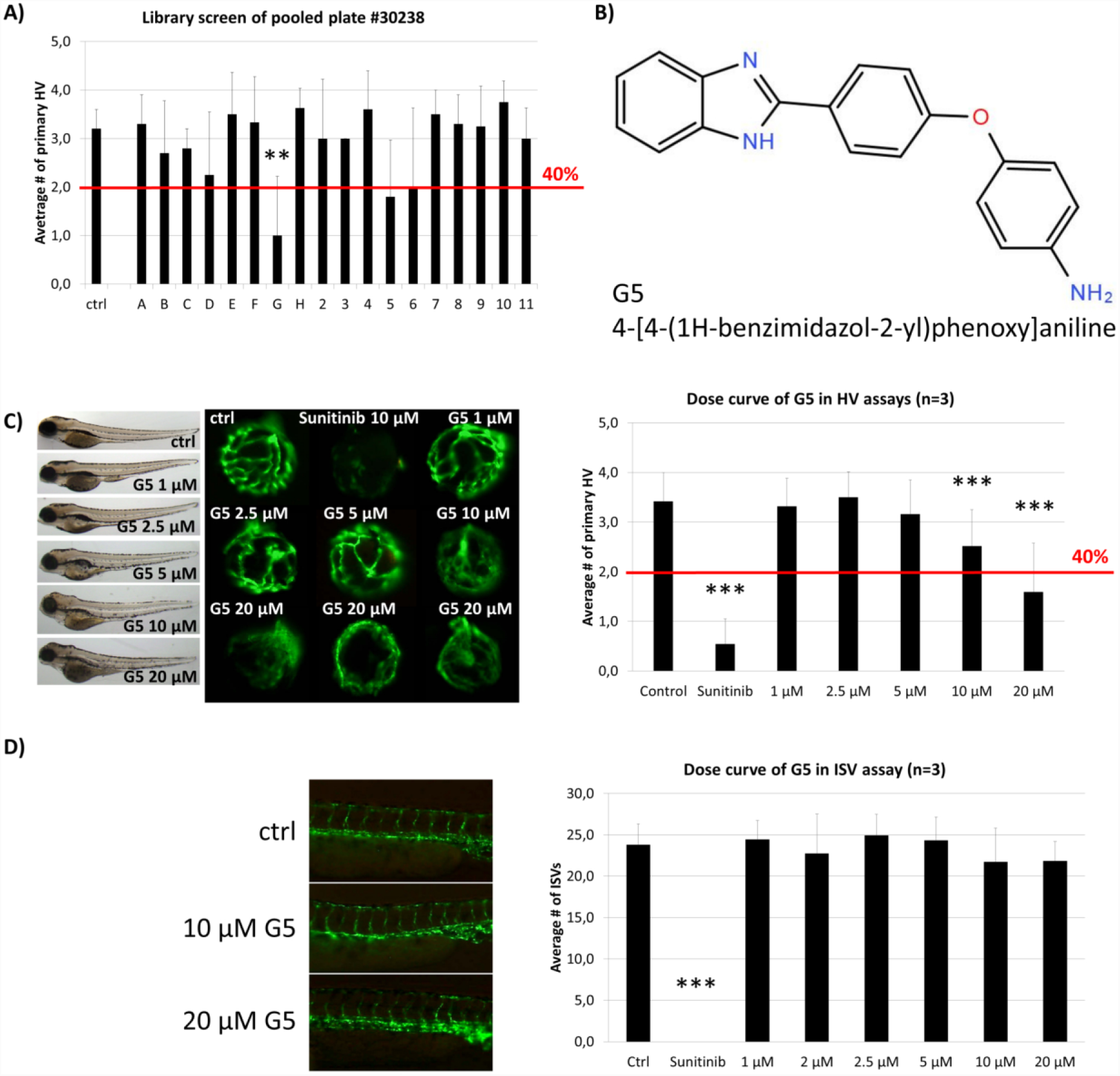
Identification and Confirmation of Hit G5. (A) G5 was a promising hit identified in the screen of DIVERSet™ library plate #30238. (B) The chemical structure and chemical name (4-[4-(1H-benzimidazol-2-yl)phenoxy]aniline) of hit G5. (C) G5 shows a dose-dependent and robust reduction of primary hyaloid vessels at 10 and 20 μM. In addition, at higher doses treated larvae show no difference in survival rates and had only mild morphological changes compared to vehicle control treatments. (D) In contrast, G5 had no significant anti-angiogenic effect in the ISV assay at the same range of concentrations. Data are means +/− SD. Statistical significance calculated by t-test and Mann Whitney test (* p < 0.05, ** p < 0.01, *** p < 0.001).

### Metrics

Our approach enabled screening of 1760 compounds of the DIVERSet™ library in only 970 treatments. A pre-screen concern with pooling was toxicity levels. We expected from our previous experience with non-pooling screens, that 2-5% of test compounds are lethal at 10 μM (Clifton et al., 2010; Merrigan and Kennedy, 2017). This predicted an average of 2-4 toxic drugs per plate which could be represented in 3-8 of the pools (expected toxicity of 17-44%). In practice, the 22% observed toxicity was lower than expected. Another pre-screen concern was the expected higher variation between technical replicates, but the Fligner-Killeen test showed no difference in variation between groups treated with one drug or a drug pool. Based on the screen metrics, the effect of the pooling strategy on sample sizes was calculated for three different libraries (100, 1,000 or 10,000 compounds) and three scenarios with the same, none or even higher toxicity as experienced in our library screen to demonstrate the usefulness of this method for other screens especially with larger libraries (***Table 2***). Notably, this pooling approach significantly reduced the required workload (***Table 1***). This screen required 534 person hours to be completed. The single most time-consuming step (50% time) was needed for the eye dissection and blood vessel quantification, which took ∼15 min per well. The remaining time was used for setting up the fish mating (∼6 min/well), collecting, sorting and distributing embryos to the wells of the plate (∼3 min/well), preparing the drug pools (∼5 min/well) and applying the treatment (∼1 min/well). In this respect, drug-pooling demonstrated its usefulness by effectively lowering sample numbers and reducing the associated time needed for analysis and not just reducing the time for screen set-up and execution as benefitted by other approaches. As a non-pooling approach would have required at least twice the number of treatments tested and analyzed, we calculate that the required time and costs incurred would have doubled if no pooling strategy was applied.

**Table 1:**
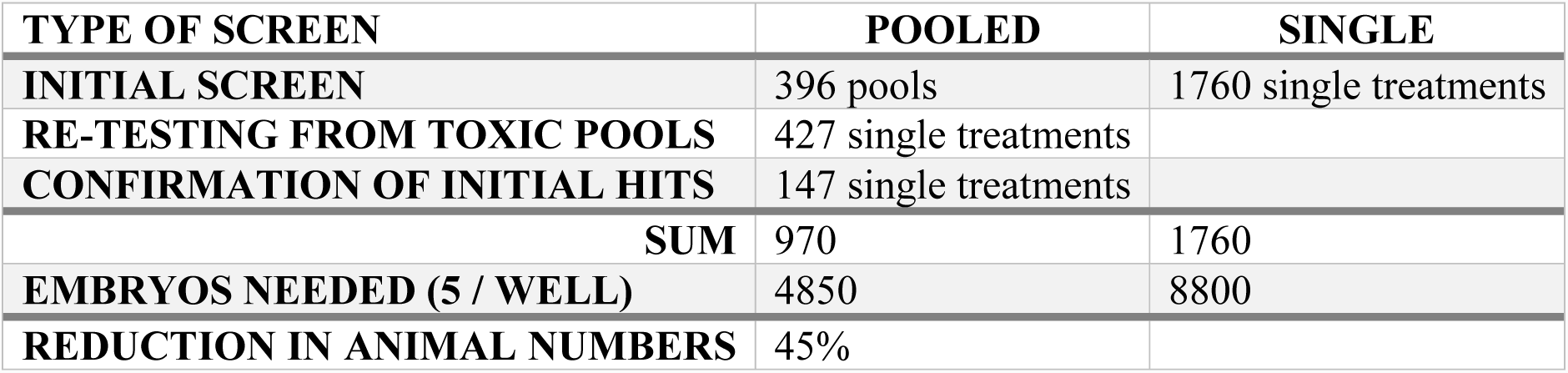
Treatment Reduction by Orthogonal Pooling. Summary of the number of immature larval zebrafish utilized for orthogonal pooled versus single “one compound, one well” approach.

**Table 2:**
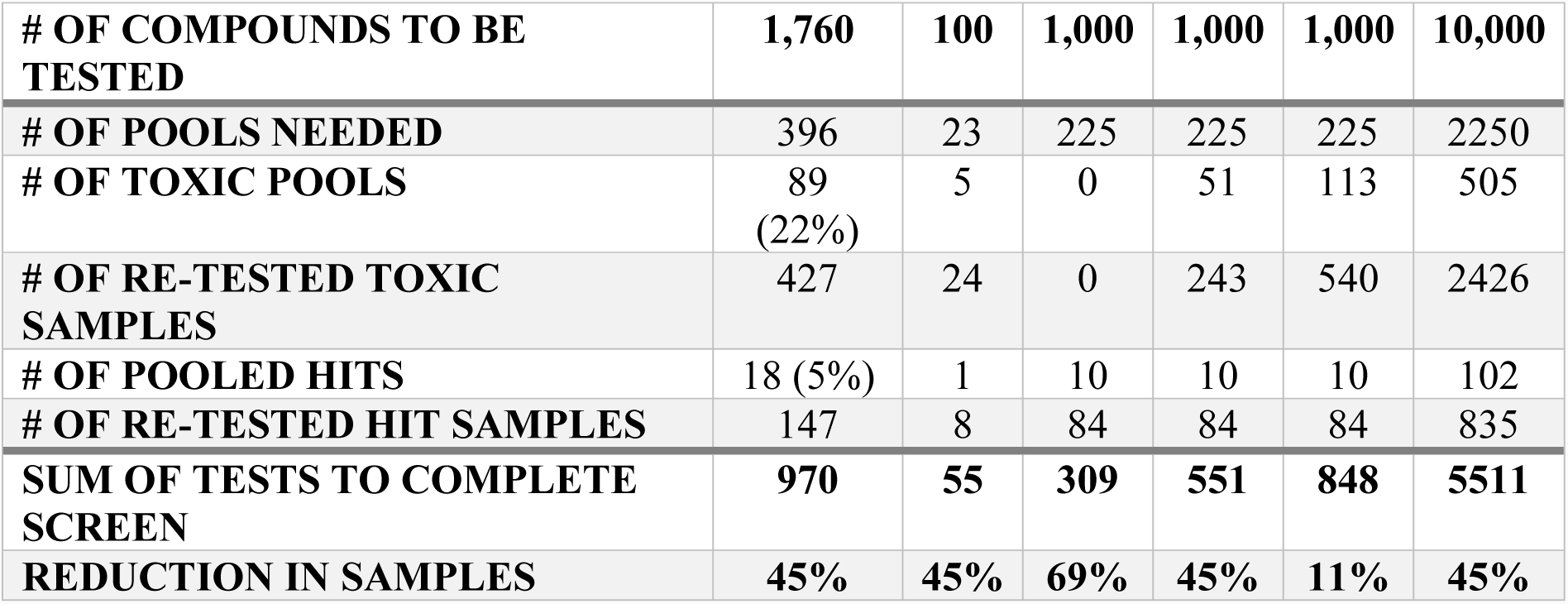
Effect of Orthogonal Pooling. Applying the screen metrics, the effect of the pooling strategy on sample size was calculated for three different sized libraries (100, 1,000 or 10,000 compounds) and with 0%, 22% and 50% toxic pools.

| # OF COMPOUNDS TO BE TESTED | 1,760 | 100 | 1,000 | 1,000 | 1,000 | 10,000 |
| --- | --- | --- | --- | --- | --- | --- |
| # OF POOLS NEEDED | 396 | 23 | 225 | 225 | 225 | 2250 |
| # OF TOXIC POOLS | 89<br>(22%) | 5 | 0 | 51 | 113 | 505 |
| # OF RE-TESTED TOXIC SAMPLES | 427 | 24 | 0 | 243 | 540 | 2426 |
| # OF POOLED HITS | 18 (5%) | 1 | 10 | 10 | 10 | 102 |
| # OF RE-TESTED HIT SAMPLES | 147 | 8 | 84 | 84 | 84 | 835 |
| SUM OF TESTS TO COMPLETE SCREEN | 970 | 55 | 309 | 551 | 848 | 5511 |
| REDUCTION IN SAMPLES | 45% | 45% | 69% | 45% | 11% | 45% |

## Discussion

Phenotype-based drug screening is still the most productive approach to discover *first-in-class* drugs (Swinney and Anthony, 2011; Zheng et al., 2013; Wagner, 2016; Moffat et al., 2017). Zebrafish is a prolific vertebrate model for forward pharmacology combining high predictive power with throughput (Peterson, 2004; Ridges et al., 2012; Rennekamp and Peterson, 2015; Williams and Hong, 2016; Ribeiro et al., 2018). With a rapidly expanding suite of transgenic, knockout and knockin lines available accelerated by gene editing technology, ever more sophisticated assay endpoints in zebrafish are applicable to phenotype-based screens (Baraban et al., 2013; Jimenez et al., 2016; Liu et al., 2016; Ribeiro et al., 2018). A number of phenotype-based screens in zebrafish, testing single drugs, discovered or validated anti-angiogenic small molecule drugs (Tran et al., 2007; Kitambi et al., 2009; Reynolds et al., 2016; Butler et al., 2017). Notably, many hits are also anti-angiogenic in mammalian models, indeed several are market authorized for clinical use in humans (Chan et al., 2002; Reynolds et al., 2016; Rezzola et al., 2016).

Our aim was to increase the efficiency of unbiased chemical screens in zebrafish. In particular, our objective was to more effectively screen a randomized chemical library to identify hits with anti-angiogenic activity in the eye. Prior experience in randomized chemical library screening estimated that less than 1% of test drugs are true hits (Reynolds et al., 2016). A “one compound, one well” approach was too time-consuming. An adaptive pooling strategy where every compound is tested in only one pool increased the risk of false negatives. Thus, our approach was to treat *Tg(fli1a:eGFP)* zebrafish larvae with 8 or 10 drugs simultaneously, applying an orthogonal pooling strategy wherein every compound is represented in two pools (Motlekar et al., 2008; Kainkaryam and Woolf, 2009; Paiva et al., 2017). Selection of this strategy was based on the following rationale: *i)* most randomized chemical library compounds are inactive in a given assay, *ii)* robust inhibitors of angiogenesis can retain activity in drug pools and *iii*) orthogonal pooling would lower the likelihood of false positive or negatives due to synergistic or antagonistic effects (Ferrand et al., 2005; Usha Warrior, 2007).

Typically, we screen drugs in the presence of 0.1% DMSO (Sasore and Kennedy, 2014; Reynolds et al., 2016; Tal et al., 2017). As the DIVERSET® compounds are provided at 10 mM stock in 100% DMSO, we confirmed at the outset that the 1% DMSO concentration in all pools did not elicit toxic/pharmacological effects on hyaloid vasculature development. In accordance with previous investigations, we didn’t observe an adverse effect of 1% DMSO on larval zebrafish development (***Fig. 2B***) (Hallare et al., 2006; Xiong et al., 2017).Overall, toxicity affected 22% of the pools representing 24% of compounds in the screen. In contrast, only 3% of single drug treated resulted in toxicity. This suggests that the toxic effects arise from the drug combinations more than individual drugs, due to additive or synergistic effects causing higher toxicity (Foucquier and Guedj, 2015). Generally, zebrafish embryos are a good model for predicting toxicity, and testing on whole organisms can prevent more costly drug failure setbacks with off-target effects later in drug development (Raldua and Pina, 2014). Despite an increased toxicity, resulting in 424 samples requiring re-testing, the sample numbers (immature larvae) required for the pooling approach was reduced by 45% compared to a “*one compound, one well*” approach (***Table 1***). Therefore, our pooling screen in immature zebrafish larvae replaces the use of animals and significantly reduces the number of immature animal forms tested, in accordance with the 3R principles (Brannen et al., 2016). For libraries with less bioactive substances or tested at lower concentrations, a theoretical sample size reduction of 69% is possible (***Table 2***). Even if twice the number of pools were impossible to analyze due to toxicity and 50% of the samples had to be re-analyzed individually, the overall sample size would reduce by 11%. Testing in drug pools were expected to cause higher variation in results from replicates. However, the statistical analysis showed no significant difference in variation of groups treated with one drug or a drug pool. Still, we concentrated our efforts on compounds with a high threshold of 40% reduction of primary hyaloid vessel numbers to be considered a hit. This design strategy was successful in screening for anti-angiogenic drugs.

In conclusion, phenotype-based drug screening in zebrafish remains a powerful approach for drug discovery. Orthogonal drug pooling strategies can easily be applied to other screening paradigms in zebrafish. Orthogonal drug pooling increases screening throughput and reduces sample numbers requirements, significantly saving time and costs and complying with the 3R principles.

## Acknowledgements

We thank members of the UCD Ocular Pharmacology and Genetics Group for comments on the draft manuscript. We thank UCD technical staff for care of the zebrafish and maintenance of the UCD Conway zebrafish facility.

## Authors Contribution

NO, TS, YA and BK contributed conception and design of the study; NO and TS performed experiments; DH and MC performed statistical analysis; NO wrote first draft of the manuscript. All authors contributed to manuscript revision, read and approved the submitted version.

## References

Alvarez, Y., Cederlund, M.L., Cottell, D.C., Bill, B.R., Ekker, S.C., Torres-Vazquez, J., et al. (2007). Genetic determinants of hyaloid and retinal vasculature in zebrafish. BMC Dev Biol 7, 114. doi: 10.1186/1471-213X-7-114.

Baraban, S.C., Dinday, M.T., and Hortopan, G.A. (2013). Drug screening in Scn1a zebrafish mutant identifies clemizole as a potential Dravet syndrome treatment. Nat Commun 4, 2410. doi: 10.1038/ncomms3410.

Bourne, R.R., Stevens, G.A., White, R.A., Smith, J.L., Flaxman, S.R., Price, H., et al. (2013). Causes of vision loss worldwide, 1990-2010: a systematic analysis. Lancet Glob Health 1(6), e339–349. doi: 10.1016/S2214-109X(13)70113-X.

Brannen, K.C., Chapin, R.E., Jacobs, A.C., and Green, M.L. (2016). Alternative Models of Developmental and Reproductive Toxicity in Pharmaceutical Risk Assessment and the 3Rs. ILAR J 57(2), 144–156. doi: 10.1093/ilar/ilw026.

Breitwieser, H., Dickmeis, T., Vogt, M., Ferg, M., and Pylatiuk, C. (2018). Fully Automated Pipetting Sorting System for Different Morphological Phenotypes of Zebrafish Embryos. SLAS Technol 23(2), 128–133. doi: 10.1177/2472630317745780.

Burns, C.G., Milan, D.J., Grande, E.J., Rottbauer, W., MacRae, C.A., and Fishman, M.C. (2005). High-throughput assay for small molecules that modulate zebrafish embryonic heart rate. Nat Chem Biol 1(5), 263–264. doi: 10.1038/nchembio732.

Butler, C.T., Reynolds, A.L., Tosetto, M., Dillon, E.T., Guiry, P.J., Cagney, G., et al. (2017). A Quininib Analogue and Cysteinyl Leukotriene Receptor Antagonist Inhibits Vascular Endothelial Growth Factor (VEGF)-independent Angiogenesis and Exerts an Additive Antiangiogenic Response with Bevacizumab. J Biol Chem 292(9), 3552–3567. doi: 10.1074/jbc.M116.747766.

Chan, J., Bayliss, P.E., Wood, J.M., and Roberts, T.M. (2002). Dissection of angiogenic signaling in zebrafish using a chemical genetic approach. Cancer Cell 1(3), 257–267.

Clifton, J.D., Lucumi, E., Myers, M.C., Napper, A., Hama, K., Farber, S.A., et al. (2010). Identification of novel inhibitors of dietary lipid absorption using zebrafish. PLoS One 5(8), e12386. doi: 10.1371/journal.pone.0012386.

Cunha-Vaz, J. (2014). Phenotypes and biomarkers of diabetic retinopathy. Personalized medicine for diabetic retinopathy: the Weisenfeld award. Invest Ophthalmol Vis Sci 55(8), 5412–5419. doi: 10.1167/iovs.14-14884.

Faivre, S., Demetri, G., Sargent, W., and Raymond, E. (2007). Molecular basis for sunitinib efficacy and future clinical development. Nat Rev Drug Discov 6(9), 734–745. doi: 10.1038/nrd2380.

Ferrand, S., Schmid, A., Engeloch, C., and Glickman, J.F. (2005). Statistical evaluation of a self-deconvoluting matrix strategy for high-throughput screening of the CXCR3 receptor. Assay Drug Dev Technol 3(4), 413–424. doi: 10.1089/adt.2005.3.413.

Foucquier, J., and Guedj, M. (2015). Analysis of drug combinations: current methodological landscape. Pharmacol Res Perspect 3(3), e00149. doi: 10.1002/prp2.149.

Goodwin, N., Westall, L., Karp, N.A., Hazlehurst, D., Kovacs, C., Keeble, R., et al. (2016). Evaluating and Optimizing Fish Health and Welfare During Experimental Procedures. Zebrafish 13 Suppl 1, S127–131. doi: 10.1089/zeb.2015.1165.

Graf, S.F., Hotzel, S., Liebel, U., Stemmer, A., and Knapp, H.F. (2011). Image-based fluidic sorting system for automated Zebrafish egg sorting into multiwell plates. J Lab Autom 16(2), 105–111. doi: 10.1016/j.jala.2010.11.002.

Hallare, A., Nagel, K., Kohler, H.R., and Triebskorn, R. (2006). Comparative embryotoxicity and proteotoxicity of three carrier solvents to zebrafish (Danio rerio) embryos. Ecotoxicol Environ Saf 63(3), 378–388. doi: 10.1016/j.ecoenv.2005.07.006.

Hao, Z., and Sadek, I. (2016). Sunitinib: the antiangiogenic effects and beyond. Onco Targets Ther 9, 5495–5505. doi: 10.2147/OTT.S112242.

Hartsock, A., Lee, C., Arnold, V., and Gross, J.M. (2014). In vivo analysis of hyaloid vasculature morphogenesis in zebrafish: A role for the lens in maturation and maintenance of the hyaloid. Dev Biol 394(2), 327–339. doi: 10.1016/j.ydbio.2014.07.024.

Isogai, S., Horiguchi, M., and Weinstein, B.M. (2001). The vascular anatomy of the developing zebrafish: an atlas of embryonic and early larval development. Dev Biol 230(2), 278–301. doi: 10.1006/dbio.2000.9995.

Jimenez, L., Wang, J., Morrison, M.A., Whatcott, C., Soh, K.K., Warner, S., et al. (2016). Phenotypic chemical screening using a zebrafish neural crest EMT reporter identifies retinoic acid as an inhibitor of epithelial morphogenesis. Dis Model Mech 9(4), 389–400. doi: 10.1242/dmm.021790.

Kainkaryam, R.M., and Woolf, P.J. (2009). Pooling in high-throughput drug screening. Curr Opin Drug Discov Devel 12(3), 339–350.

Kitambi, S.S., McCulloch, K.J., Peterson, R.T., and Malicki, J.J. (2009). Small molecule screen for compounds that affect vascular development in the zebrafish retina. Mech Dev 126(5-6), 464–477.

Kwong, T.Q., and Mohamed, M. (2014). Anti-vascular endothelial growth factor therapies in ophthalmology: current use, controversies and the future. Br J Clin Pharmacol 78(4), 699–706. doi: 10.1111/bcp.12371.

Lawson, N.D., and Weinstein, B.M. (2002). In vivo imaging of embryonic vascular development using transgenic zebrafish. Dev Biol 248(2), 307–318.

Li, P., Lahvic, J.L., Binder, V., Pugach, E.K., Riley, E.B., Tamplin, O.J., et al. (2015). Epoxyeicosatrienoic acids enhance embryonic haematopoiesis and adult marrow engraftment. Nature 523(7561), 468–471. doi: 10.1038/nature14569.

Liu, H., Chen, S., Huang, K., Kim, J., Mo, H., Iovine, R., et al. (2016). A High-Content Larval Zebrafish Brain Imaging Method for Small Molecule Drug Discovery. PLoS One 11(10), e0164645. doi: 10.1371/journal.pone.0164645.

MacRae, C.A., and Peterson, R.T. (2015). Zebrafish as tools for drug discovery. Nat Rev Drug Discov 14(10), 721–731. doi: 10.1038/nrd4627.

Merrigan, S.L., and Kennedy, B.N. (2017). Vitamin D receptor agonists regulate ocular developmental angiogenesis and modulate expression of dre-miR-21 and VEGF. Br J Pharmacol 174(16), 2636–2651. doi: 10.1111/bph.13875.

Moffat, J.G., Vincent, F., Lee, J.A., Eder, J., and Prunotto, M. (2017). Opportunities and challenges in phenotypic drug discovery: an industry perspective. Nat Rev Drug Discov 16(8), 531–543. doi: 10.1038/nrd.2017.111.

Motlekar, N., Diamond, S.L., and Napper, A.D. (2008). Evaluation of an orthogonal pooling strategy for rapid high-throughput screening of proteases. Assay Drug Dev Technol 6(3), 395–405. doi: 10.1089/adt.2007.110.

North, T.E., Goessling, W., Walkley, C.R., Lengerke, C., Kopani, K.R., Lord, A.M., et al. (2007). Prostaglandin E2 regulates vertebrate haematopoietic stem cell homeostasis. Nature 447(7147), 1007–1011. doi: 10.1038/nature05883.

Paiva, A.A., Klakouski, C., Li, S., Johnson, B.M., Shu, Y.Z., Josephs, J., et al. (2017). Development, optimization and implementation of a centralized metabolic soft spot assay. Bioanalysis 9(7), 541–552. doi: 10.4155/bio-2016-0299.

Peal, D.S., Mills, R.W., Lynch, S.N., Mosley, J.M., Lim, E., Ellinor, P.T., et al. (2011). Novel chemical suppressors of long QT syndrome identified by an in vivo functional screen. Circulation 123(1), 23–30. doi: 10.1161/CIRCULATIONAHA.110.003731.

Peterson, R.T. (2004). Discovery of therapeutic targets by phenotype-based zebrafish screens. Drug Discov Today Technol 1(1), 49–54. doi: 10.1016/j.ddtec.2004.07.002.

Peterson, R.T., Link, B.A., Dowling, J.E., and Schreiber, S.L. (2000). Small molecule developmental screens reveal the logic and timing of vertebrate development. Proc Natl Acad Sci U S A 97(24), 12965–12969. doi: 10.1073/pnas.97.24.12965.

Querques, G., Capuano, V., Frascio, P., Bandello, F., and Souied, E.H. (2015). Emerging therapeutic options in age-related macular degeneration. Ophthalmic Res 53(4), 194–199. doi: 10.1159/000379754.

Rai, A.K., and Sherkow, J.S. (2016). The changing life science patent landscape. Nat Biotechnol 34(3), 292–294. doi: 10.1038/nbt.3504.

Raldua, D., and Pina, B. (2014). In vivo zebrafish assays for analyzing drug toxicity. Expert Opin Drug Metab Toxicol 10(5), 685–697. doi: 10.1517/17425255.2014.896339.

Rennekamp, A.J., Huang, X.P., Wang, Y., Patel, S., Lorello, P.J., Cade, L., et al. (2016). sigma1 receptor ligands control a switch between passive and active threat responses. Nat Chem Biol 12(7), 552–558. doi: 10.1038/nchembio.2089.

Rennekamp, A.J., and Peterson, R.T. (2015). 15 years of zebrafish chemical screening. Curr Opin Chem Biol 24, 58–70. doi: 10.1016/j.cbpa.2014.10.025.

Reynolds, A.L., Alvarez, Y., Sasore, T., Waghorne, N., Butler, C.T., Kilty, C., et al. (2016). Phenotype-based Discovery of 2-[(E)-2-(Quinolin-2-yl)vinyl]phenol as a Novel Regulator of Ocular Angiogenesis. J Biol Chem 291(14), 7242–7255. doi: 10.1074/jbc.M115.710665.

Rezzola, S., Paganini, G., Semeraro, F., Presta, M., and Tobia, C. (2016). Zebrafish (Danio rerio) embryo as a platform for the identification of novel angiogenesis inhibitors of retinal vascular diseases. Biochim Biophys Acta 1862(7), 1291–1296. doi: 10.1016/j.bbadis.2016.04.009.

Ribeiro, C.J.A., Kankanala, J., Shi, K., Kurahashi, K., Kiselev, E., Ravji, A., et al. (2018). New fluorescence-based high-throughput screening assay for small molecule inhibitors of tyrosyl-DNA phosphodiesterase 2 (TDP2). Eur J Pharm Sci 118, 67–79. doi: 10.1016/j.ejps.2018.03.021.

Ridges, S., Heaton, W.L., Joshi, D., Choi, H., Eiring, A., Batchelor, L., et al. (2012). Zebrafish screen identifies novel compound with selective toxicity against leukemia. Blood 119(24), 5621–5631. doi: 10.1182/blood-2011-12-398818.

Sasore, T., and Kennedy, B. (2014). Deciphering combinations of PI3K/AKT/mTOR pathway drugs augmenting anti-angiogenic efficacy in vivo. PLoS One 9(8), e105280. doi: 10.1371/journal.pone.0105280.

Saydmohammed, M., Vollmer, L.L., Onuoha, E.O., Maskrey, T.S., Gibson, G., Watkins, S.C., et al. (2018). A High-Content Screen Reveals New Small-Molecule Enhancers of Ras/Mapk Signaling as Probes for Zebrafish Heart Development. Molecules 23(7). doi: 10.3390/molecules23071691.

Scannell, J.W., Blanckley, A., Boldon, H., and Warrington, B. (2012). Diagnosing the decline in pharmaceutical R&D efficiency. Nat Rev Drug Discov 11(3), 191–200. doi: 10.1038/nrd3681.

Swinney, D.C., and Anthony, J. (2011). How were new medicines discovered? Nat Rev Drug Discov 10(7), 507–519. doi: 10.1038/nrd3480.

Tal, T., Kilty, C., Smith, A., LaLone, C., Kennedy, B., Tennant, A., et al. (2017). Screening for angiogenic inhibitors in zebrafish to evaluate a predictive model for developmental vascular toxicity. Reprod Toxicol 70, 70–81. doi: 10.1016/j.reprotox.2016.12.004.

Tran, T.C., Sneed, B., Haider, J., Blavo, D., White, A., Aiyejorun, T., et al. (2007). Automated, quantitative screening assay for antiangiogenic compounds using transgenic zebrafish. Cancer Res 67(23), 11386–11392. doi: 10.1158/0008-5472.CAN-07-3126.

Usha Warrior, S.M.G., Linda Traphagen, Gail Freiberg, Danli Towne, Patrick Humphrey, James Kofron, David J. Burns. (2007). Maximizing the Identification of Leads from Compound Mixtures. Letters in Drug Design & Discovery 4(3), 215 – 223. doi: 10.2174/157018007780077408.

Vogt, A., Cholewinski, A., Shen, X., Nelson, S.G., Lazo, J.S., Tsang, M., et al. (2009). Automated image-based phenotypic analysis in zebrafish embryos. Dev Dyn 238(3), 656–663. doi: 10.1002/dvdy.21892.

Wagner, B.K. (2016). The resurgence of phenotypic screening in drug discovery and development. Expert Opin Drug Discov 11(2), 121–125. doi: 10.1517/17460441.2016.1122589.

Wang, C., Tao, W., Wang, Y., Bikow, J., Lu, B., Keating, A., et al. (2010). Rosuvastatin, identified from a zebrafish chemical genetic screen for antiangiogenic compounds, suppresses the growth of prostate cancer. Eur Urol 58(3), 418–426. doi: 10.1016/j.eururo.2010.05.024.

Westerfield, M. (1995). The Zebrafish Book: A Guide for the Laboratory Use of Zebrafish (Brachydanio rerio). University of Oregon Press.

Wheeler, G.N., and Brandli, A.W. (2009). Simple vertebrate models for chemical genetics and drug discovery screens: lessons from zebrafish and Xenopus. Dev Dyn 238(6), 1287–1308. doi: 10.1002/dvdy.21967.

Wiley, D.S., Redfield, S.E., and Zon, L.I. (2017). Chemical screening in zebrafish for novel biological and therapeutic discovery. Methods Cell Biol 138, 651–679. doi: 10.1016/bs.mcb.2016.10.004.

Williams, C.H., and Hong, C.C. (2016). Zebrafish small molecule screens: Taking the phenotypic plunge. Comput Struct Biotechnol J 14, 350–356. doi: 10.1016/j.csbj.2016.09.001.

Xiong, X., Luo, S., Wu, B., and Wang, J. (2017). Comparative Developmental Toxicity and Stress Protein Responses of Dimethyl Sulfoxide to Rare Minnow and Zebrafish Embryos/Larvae. Zebrafish 14(1), 60–68. doi: 10.1089/zeb.2016.1287.

Zheng, W., Thorne, N., and McKew, J.C. (2013). Phenotypic screens as a renewed approach for drug discovery. Drug Discov Today 18(21-22), 1067–1073. doi: 10.1016/j.drudis.2013.07.001.

